# Drug repurposing of dopaminergic drugs to inhibit Ataxin-3 aggregation

**DOI:** 10.1101/2022.12.01.518659

**Authors:** Francisco Figueiredo, Zsuzsa Sárkány, Alexandra Silva, Pedro M. Martins, Sandra Macedo Ribeiro

**Affiliations:** Instituto de Investigação e Inovação em Saúde (i3S), Universidade do Porto, Porto, Portugal; Instituto de Biologia Molecular e Celular (IBMC), Universidade do Porto, Porto, Portugal; Instituto de Ciências Biomédicas Abel Salazar (ICBAS), Universidade do Porto, Porto, Portugal

## Abstract

The accumulation of mutant ataxin-3 (Atx3) in neuronal nuclear inclusions is a pathological hallmark of Machado-Joseph disease (MJD), also known as Spinocerebellar Ataxia Type 3. Decreasing the protein aggregation burden is a possible disease-modifying strategy to tackle MJD and other neurodegenerative disorders for which only symptomatic treatments are currently available. We performed a drug repurposing screening to identify inhibitors of Atx3 aggregation with known toxicological and pharmacokinetic profiles. Interestingly, dopamine hydrochloride and other catecholamines are among the most potent inhibitors of Atx3 aggregation *in vitro*. Our results indicate that low micromolar concentrations of dopamine markedly delay the formation of mature amyloid fibrils of mutant Atx3 through the inhibition of the earlier oligomerization steps. Although dopamine itself does not pass the blood-brain barrier, dopamine levels in the brain can be increased by low doses of dopamine precursors and dopamine agonists commonly used to treat Parkinsonian symptoms. These findings disclose a possible application of dopaminergic drugs to halt or reduce Atx3 accumulation in the brains of MJD patients.

## Introduction

Machado-Joseph disease (MJD), also known as Spinocerebellar Ataxia Type 3 is an inherited autosomal dominant neurodegenerative disorder. MJD is a progressive disorder of late-onset, caused by the expansion of an unstable CAG repeat in the coding region of the *ATXN3* gene, which is translated into an expanded polyglutamine (polyQ) tract in the protein ataxin-3 (Atx3). The affected brain regions are typically the dentate nucleus and the substantia nigra, although the involvement of other central nervous system regions is also reported [1]. Neuronal nuclear inclusions (NNI) rich in the expanded protein are identified selectively in the degenerating regions of the brain [2–4]. Normal and expanded Atx3 are widely expressed in the brain and throughout the body [5], and both are present in the NNI together with several other proteins such as ubiquitin and components of the cellular proteostasis machinery [6,7]. With currently no available disease-modifying therapies for MJD, several symptomatic therapies can be prescribed based on the knowledge of other related diseases and the patient’s needs [8]. For example, dopaminergic drugs such as levodopa and dopamine agonists can be used to ameliorate the parkinsonian symptoms that characterize a MJD subtype associated with CAG expansions within the 60-70 repeat ranges [9–11].

Although the precise role of NNI in MJD pathogenesis is not clear [12–14], suppression of polyQ aggregation is an attractive therapeutic strategy to tackle this and other types of polyglutamine expansion disorders [15]. For that, different indirect approaches are conceived involving, for example, modulation of the proteasome activity, autophagy induction, transcriptional regulation, and the inhibition of calcium-dependent, cysteine proteases [8,16]. Toxic gain-of-function of mutant Atx3 can be directly targeted by lowering the protein levels using antisense oligonucleotides. The positive effects of lowering the expression of polyQ-expanded Atx3 in cell models and transgenic rodents were demonstrated by several authors [17–19], thus providing evidence in support of the ongoing pharmacokinetic and safety studies of antisense oligonucleotides in MJD (ClinicalTrials.gov Identifier: NCT05160558). Ideally, the safe and effective inhibition of Atx3 aggregation should abolish the formation of NNI known as major neuropathological hallmarks of MJD without compromising the still incompletely disclosed cellular functions of Atx3.

The mechanism of Atx3 aggregation that underlies NNI formation comprises a polyQ-independent step originating SDS-sensitive protofibrils, followed by a polyQ-dependent step that originates SDS-resistant mature fibrils and is only observed for Atx3 with pathological polyQ tract sizes [20]. The first step is mediated by the Josephin domain of Atx3 and can be followed over time using the amyloid-specific fluorescent dye thioflavin-T (ThT) [21–23]. In parallel with this amyloidogenic pathway, Atx3 does also occur in the form of soluble oligomers that eventually dissociate into monomers to feed the formation of the more stable fibrillar aggregates [24]. Pre-existing fibrils of Atx3 autocatalytically promote the amyloidogenic pathway through the secondary nucleation of new aggregates and, to a lesser extent, fibril elongation [25]. Translating new drug candidates to clinical trials requires robust preclinical studies showing the safety and efficacy of the new compounds but also large cohorts of patients and long periods of observation [8,26]. The cost and duration of this process would be drastically reduced if drugs already approved for the treatment of other diseases could be repurposed for MJD therapy. Based on a ThT fluorescence assay recently optimized by us [21], a chemical library of drug repurposing compounds was screened for candidate inhibitors of Atx3 aggregation. Hit validation was performed using a combination of Transmission Electron Microscopy (TEM), Size Exclusion Chromatography (SEC), and Dynamic Light Scattering (DLS) techniques. The presence of dopamine and other catecholamines related to dopamine in the list of the most potent inhibitors prompts us to envisage a novel use of dopaminergic drugs to reduce Atx3 aggregation and prevent NNI accumulation in the brains of MJD patients.

## Materials and Methods

### Protein Expression and Purification

Both Atx3 isoforms were expressed and purified as previously described [21]. After SDS-PAGE analysis, fractions corresponding to Atx3 were pooled and concentrated on an Amicon Ultra-15 centrifugal filter (10 kDa MWCO, Millipore) to 20-30 mg.mL^-1^ in buffer A (20 mM sodium phosphate pH 7.5, 150 mM NaCl, 5% (v/v) glycerol, 2 mM EDTA, 1 mM DTT) frozen in liquid nitrogen, and stored at −80 °C. Before each aggregation assay, purified Atx3 aliquots were applied to a Superose 12 10/300 GL column (GE Healthcare Life Sciences) pre-equilibrated with aggregation buffer (20 mM HEPES pH 7.5, 150 mM NaCl, and 1 mM DTT). The final protein concentration was determined by measuring the absorbance at 280 nM and using the molar extinction coefficient (31650 M^-1^.cm^-1^).

### Thioflavin-T Aggregation Assay

Atx3 aggregation was monitored by following the increase in thioflavin T (ThT) fluorescence at 480 nm (440 nm excitation) on a CHAMELEON V plate reader (HIDEX) using a 384-well microplate (low flange, black, flat bottom, polystyrene; Corning, Kennebunk, ME). Briefly, 50 μL samples of Atx3 at 3 μM in aggregation buffer containing 15 μM ThT were incubated at 37 °C and ThT fluorescence was measured every 30 min for approximately 48h. To prevent evaporation, 20 μL of paraffin oil was added in each reaction well with an automatic multichannel pipette (Eppendorf Xplorer, ref 4861000120).

### Compound Screening

The commercial chemical library (Prestwick, Illkirch) containing 1280 drug repurposing compounds was screened to identify Atx3 aggregation inhibitors. Atx3 with a homorepeat segment containing 13 glutamines (Atx3 13Q) was dispensed into a 384-well microplate with an automatic multichannel pipette (Eppendorf Xplorer, ref 4861000120, Eppendorf, Germany). Then an automated liquid handler (JANUS Automated Workstation; PerkinElmer) equipped with pin tool replicators (V&P Scientific) coupled to a Modular Dispense Technology (MDT) head was used to add 0.1 μL of test compounds (from a 1 mM stock) or DMSO (for controls, final concentration 0.02 % (v/v)). A total of four 384-well microplates were filled, each plate testing 320 chemical compounds (1 compound per well) and running 32 control reactions in the presence of DMSO. The final reaction mixture contained 3 μM of Atx3 13Q and 2 μM of test compound/DMSO.

### Dose Response Studies

Dose-response studies using the ThT aggregation assay were performed for the hit compounds selected in the screening with Atx3 isoforms containing a homorepeat sequence with 13 or 77 glutamines (Atx3 13Q or Atx3 77Q) using a protein concentration of 3 μM. Stock solutions of each compound were prepared at 40 μM and a series of serial-diluted compound concentrations were made to obtain the 12 different tested concentrations: 20, 10, 5, 2.5, 1.25, 0.63, 0.31, 0.16, 0.08, 0.04, 0.02 and 0.01 μM. Experiments for each condition were run in triplicate. Due to the limited solubility of pentetic acid in DMSO, a stock solution of 10 mM pentetic acid was prepared in 0.05 M NaOH for the titration of this compound. For the control experiments, the same scheme was adopted for both solvents used, varying the DMSO concentrations between 0.0004 % and 0.4 % (v/v), and the NaOH concentrations between 25 nM and 250 μM. All serial dilutions were prepared using an automatic multichannel pipette (Eppendorf Xplorer, ref 4861000120, Eppendorf, Germany). To determine the half-maximal inhibitory concentration (IC_50_), each progress curve was fitted to a logistic-type equation to determine the duration of the lag phase (*t*_lag_) and the maximal aggregation rate (*ν*_max_):

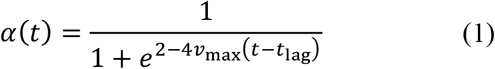

where *α*(*t*) is the reaction conversion as a function of time *t*. The fitted parameters *t*_lag_ and *ν*_max_ are related to the kinetic constants for nucleation and growth/secondary nucleation [25,27]. A four-parameter Hill equation was fitted to the concentration vs. *t*_lag_ response curve to extract IC_50_ values.

### Transmission Electron Microscopy

For visualization of protein aggregates by TEM protein samples collected at different time points of Atx3 aggregation assay were diluted 5-fold in water and adsorbed onto glow-discharged, carbon-coated films supported on 300-mesh nickel grids and negatively stained with 1 % (w/v) uranyl acetate. Grids were visualized using a JEM-1400 (JEOL, Tokyo, Japan) TEM at an accelerating voltage of 80 kV.

### Atx3 Oligomerization Analysis by Size Exclusion Chromatography

Atx3 samples (1 mL at 3 μM) were incubated in aggregation buffer without ThT for 96 hours at 37 °C without shaking. At different time points (approximately 0, 24, 48 and 72 hours), 100 μL aliquots were collected, filtered and injected in a Superdex 200 Increase 5/150 GL column (GE Healthcare Life Sciences) using a 50 μL loop. A concentration of 0.04% (v/v) DMSO or 25μM NaOH was used for the control experiments.

### Dynamic Light Scattering

Samples of 3 μM Atx3 (200 μL) in aggregation buffer were incubated in a UV-Cuvette micro cuvette (Brand, Germany) with a cap, maintained at a constant temperature of 37 °C without shaking. At different time points (0, 24, 48, 72 and 96 hours) samples were taken to Zetasizer Nano ZS DLS system (Malvern Instruments) for size distribution measurements. At least three independent measurements were made for each sample at 25 °C. A concentration of 0.04% (v/v) DMSO or 25μM NaOH was used for the control experiments. The intensity size distribution obtained by dynamic light scattering is a plot of the relative intensity of light scattered by particles in different size classes. Dynamic light scattering software (nano, Malvern Instruments) was used to determine intensity- and volume-based size distributions, sample polydispersity, and the mean hydrodynamic radius values.

### Thermal Shift Assay

The melting temperature of Atx3 in the absence and presence of the different compounds was determined using the hydrophobic fluorescent dye SYPRO Orange. For each condition, 12.5 μL of SYPRO Orange 5000x (Invitrogen) diluted to 10x in Atx3 purification buffer were mixed with 12.5 μL of ataxin-3 at (0.8 mg mL^-1^) and loaded in a white 96-well PCR plate, ref. 04-082-9056 (NerbePlus). For all the conditions three replicates were prepared. The thermal shift assay was performed in an iCycler iQ5 Multicolor Real-Time PCR detection system (Bio-Rad) running the following protocol: heating from 25 °C to 85 °C with a 30s hold time every 0.5 °C, followed by a fluorescence reading using Cy3 dye filter (excitation/emission, 545/585). Melting curves were analysed using the CFX Manager software (Bio-Rad) to calculate melting temperature (Tm) from the maximum value of the first derivative curve of the melting curve.

## Results

### Approved drugs can prevent Ataxin-3 aggregation

By employing a ThT fluorescence assay recently described and validated by us [21], 1280 compounds were screened for potential inhibitors of Atx3 aggregation. The initial identification of hit compounds was performed using the Atx3 13Q isoform because ThT fluorescence monitors steps of Atx3 aggregation that are common to both Atx3 13Q and Atx3 77Q (Figure 1A). We were particularly interested in kinetic inhibitors of the primary (1ry) and secondary (2ry) nucleation steps that are more relevant for prion-like mechanisms of disease spreading in the brain and, in the case of Atx3 aggregation, are known to dominate over the fibril elongation step [25]. Although more sophisticated analyses are required for identifying other mechanisms of inhibition, a two-point assay monitoring the ThT fluorescence increase (Δ*F*) at the end of a long incubation time (*t*\*) provides sufficient information for distinguishing kinetic inhibitors from false positives resulting from the occurrence of fluorescence artifacts [28]. Our strategy additionally contemplates kinetic measurables such as the duration of the lag phase *t*_lag_ and the maximal aggregation rate *ν*_max_ (Figure 1B) that are investigated during subsequent dose-response studies. A total of 23 compounds exhibited less than 20% of the control fluorescence increase measured at the end of *t*\* = 30 h (Figure 1C). Included in this list are catechols and catecholamines such as carbidopa, entacapone, isoproterenol, benserazide hydrochloride, levonordefrin, isoetharine mesylate salt, Isoproterenol hydrochloride, and dopamine hydrochloride, whose inhibitory effect on the aggregation of both isoforms of Atx3 was confirmed by dose-response analysis (not shown here except for dopamine). Of these compounds, only dopamine hydrochloride was selected for complementary validation analysis since pharmacological evidence is already available showing that dopamine levels in the brain can be safely increased through the use of dopaminergic drugs.

**Figure 1.**
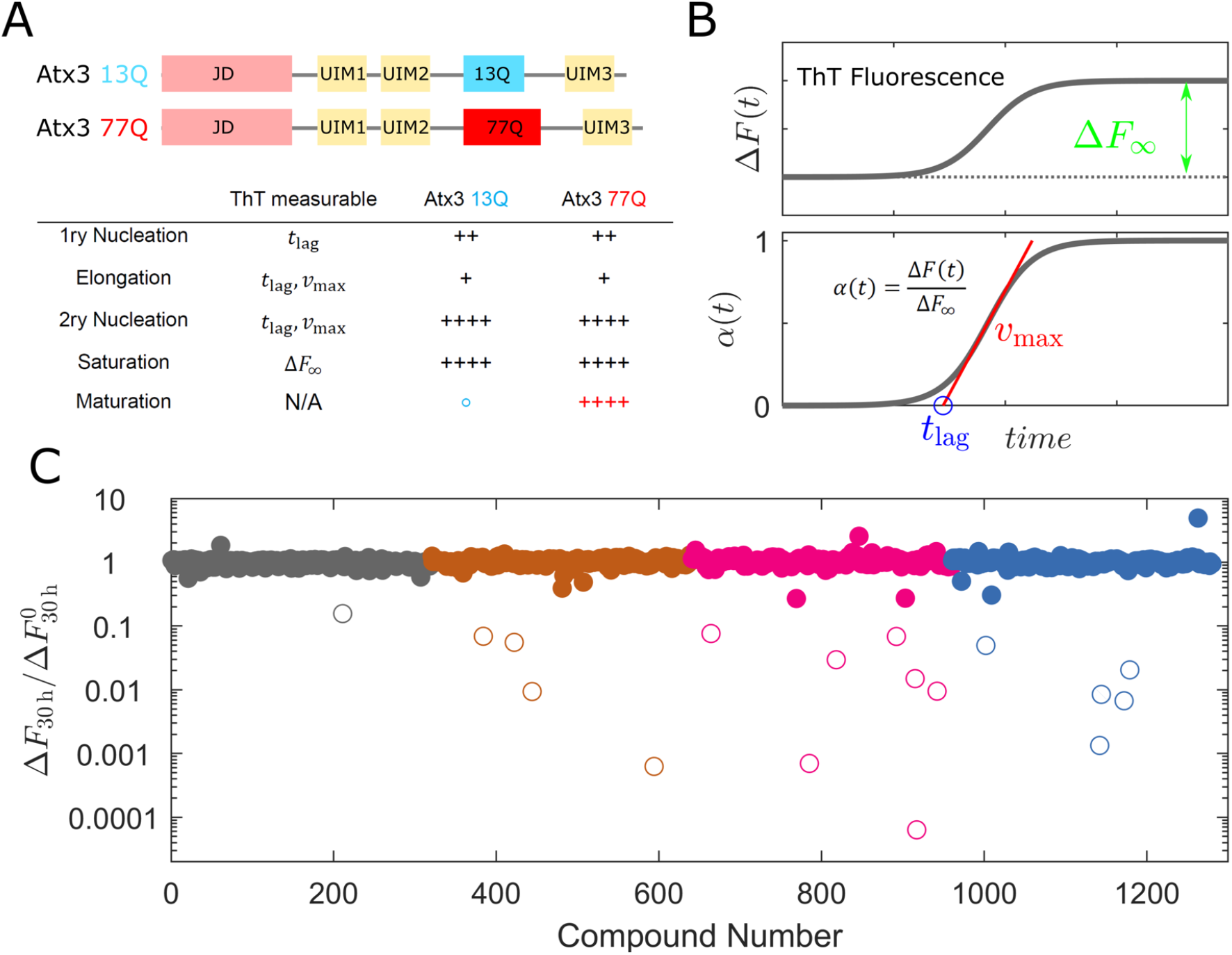
Screening strategy and first hit compounds. (A) Top: domain architectures of Atx3 13Q and Atx3 77Q. Bottom: the kinetic steps of Atx3 aggregation detected by ThT fluorescence measurables are common to both isoforms; only the expanded isoform undergoes fibril maturation. (B) Illustration of the ThT fluorescence measurables obtained from nonnormalized (top) and normalized (bottom) progress curves. (C) Symbols: Fluorescence increase measured at the end of 30 h for each tested compound over the average Δ*F* value obtained for controls 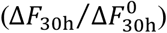. Hit compounds (open symbols) exhibit values of this ratio below 0.2. Different colors indicate the four 384-well microplates used to screen 1280 drug repurposing compounds in Atx3 13Q aggregation monitored by the increase in ThT fluorescence.

### Selection of the most potent Ataxin-3 aggregation inhibitors

To select the most potent inhibitors of Atx3 aggregation, the effect of compound concentration on the protein aggregation curves of Atx3 was measured and analyzed to extract kinetic measurables. We confirmed that dopamine hydrochloride, tolcapone, ciclopirox ethanolamine and pentetic acid strongly inhibited the aggregation of both Atx3 isoforms in a concentrationdependent manner (Figure 2).

**Figure 2.**
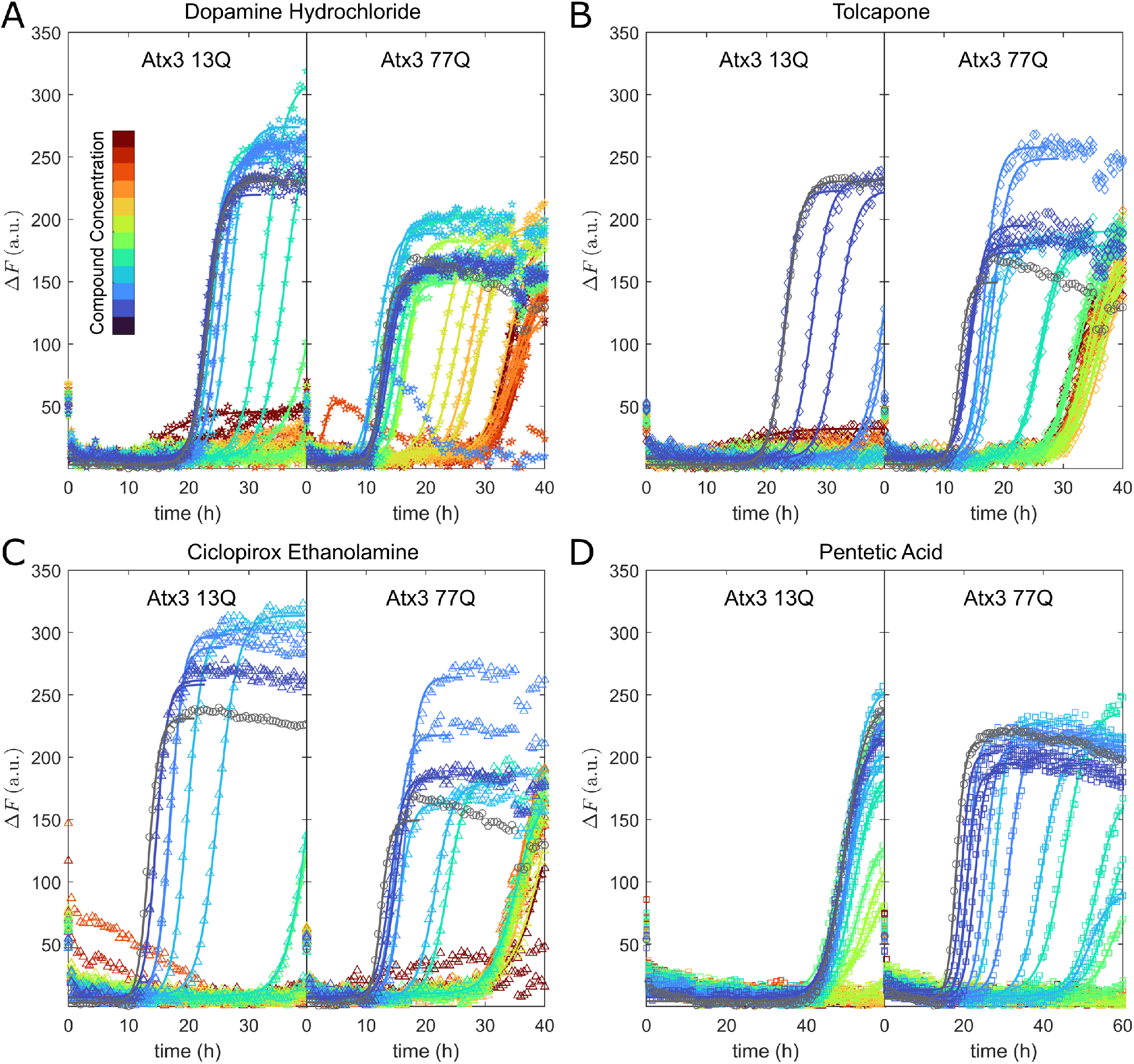
Dose-response assessment for the selected compounds. ThT fluorescence increase was measured during the aggregation of Atx3 in the presence of different concentrations of (A) dopamine hydrochloride, (B) tolcapone, (C) ciclopirox ethanolamine and (D) pentetic acid. In each panel, the left- and right-side graphs refer to Atx3 13Q and Atx3 77Q aggregation, respectively. Colors: serial dilutions from 20 μM (dark red) to 0.01 μM (dark blue) following the color sequence indicated in (A). Gray circles: average data obtained during 12 control experiments. Other symbols: Δ*F* data obtained during triplicate experiments. Lines: numerical fit by a nucleation-and-growth model.

All curves showing a meaningful increase of ThT fluorescence over time were well fit by a sigmoidal equation containing two kinetic parameters plus the upper and lower fluorescence limits (Eq. 1). The fitted rate constants were used to obtain *t*_lag_ (Figure 3A) and %max values, but only *t*_lag_ was used to estimate IC_50_ values, given that *ν*_max_ in sigmoidal progress curves is poorly affected by the 1ry nucleation step [25,27]. Submicromolar concentrations of the selected compounds effectively inhibit the aggregation of both isoforms of Atx3 (Figure 3B), although the inhibitory effect is generally more pronounced during Atx3 13Q aggregation. The longer periods required for Atx3 13Q aggregation both in the presence and absence of pentetic acid are due to the presence of NaOH, required to solubilize this compound. A reciprocal dependence of !_lag_ on *ν*_max_ is identified for all selected compounds except for the said case of Atx3 13Q aggregation in the presence of pentetic acid (Figure 3C), where longer periods of observation would be required to ascertain the *t*_lag_ vs. *ν*_max_ relationship. From this preliminary analysis, we conclude that the aggregation-delay effect induced by the compounds is the result of slower rates of secondary processes such as fibril elongation and/or 2ry nucleation. If only primary nucleation was inhibited, *ν*_max_ would not be affected by increasing concentrations of the tested compound (Figure 1A).

**Figure 3.**
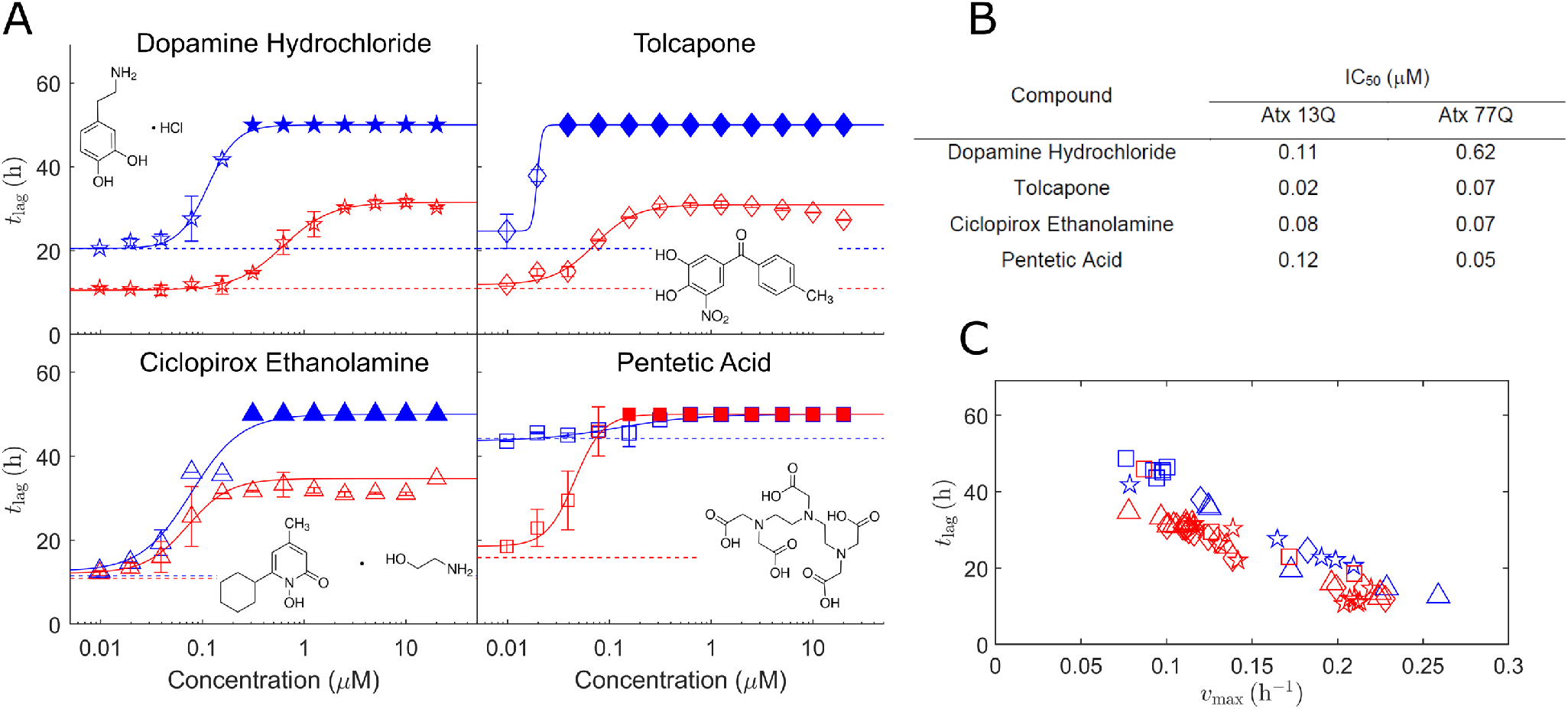
Dose-response analysis. (A) Variation of *t*_lag_ with compound concentration during the aggregation of Atx3 13Q (blue) and Atx3 77 (red). Open symbols and error bars: mean and standard deviation of triplicate measurements of *t*_lag_. Filled symbols: values of *t*_lag_ = 50 h are assumed whenever the duration of the lag phase exceeded the limit of detection of the assay. Lines: numerical fit by a concentration-response curve. Structures of each selected compound are represented in each plot. (B) Fitted values of IC_50_. (C) Relationship between mean values of *t*_lag_ and *ν*_max_ obtained in the presence of the selected compounds. Colors and symbols as in (A). A reciprocal *t*_lag_ vs. *ν*_max_ dependence indicates that mainly secondary processes (fibril elongation and/or 2ry nucleation) are targeted by the inhibitors.

### Dopamine delays the secondary nucleation of new Atx3 fibrils

Among the selected compounds, dopamine hydrochloride and, more generally, dopaminergic drugs, are particularly attractive to be repurposed as disease-modifying drugs delaying NNI accumulation. Given the preventive component of the aggregation-inhibition strategy, inhibitor administration/reinforcement should reach the affected regions of the brain and be well tolerated by MJD gene carriers on a chronic treatment basis. Pentetic acid, for example, is not blood-brain barrier permeable [29], while tolcapone and ciclopirox ethanolamine currently require more pharmacological studies before long-term administration regimes can be warranted [30–32]. In order to decipher relevant differences between the mechanisms of inhibition elicited by dopamine and the other 3 compounds, we started by using a thermal-shift assay to measure the effect of each inhibitor on the melting temperatures of Atx3 13Q and Atx3 77Q. No thermal shift was induced by any of the compounds in both Atx3 isoforms (Supplementary Figure S1), suggesting that the inhibitory effect on protein aggregation is not due to high-affinity interactions with Atx3 [33].

Next, we used DLS to investigate how the dynamic populations of monomer, primary fibrils, and secondary fibrils of Atx3 13Q are affected by each selected compound. In the presence of 5 μM dopamine hydrochloride, monomer depletion and the concomitant emergence of larger Atx3 species are visibly delayed: Atx3 13Q aggregation in the control and test compound experiments appears to be concluded after 48 hours (Figures 4A and 4B) and 96 hours (Figures 4C and 4D), respectively. Additionally, the final fibril size indicated by the intensity distributions increases from ~35 nm (Figure 5A) to ~75 nm (Figure 4C) when dopamine is present.

**Figure 4.**
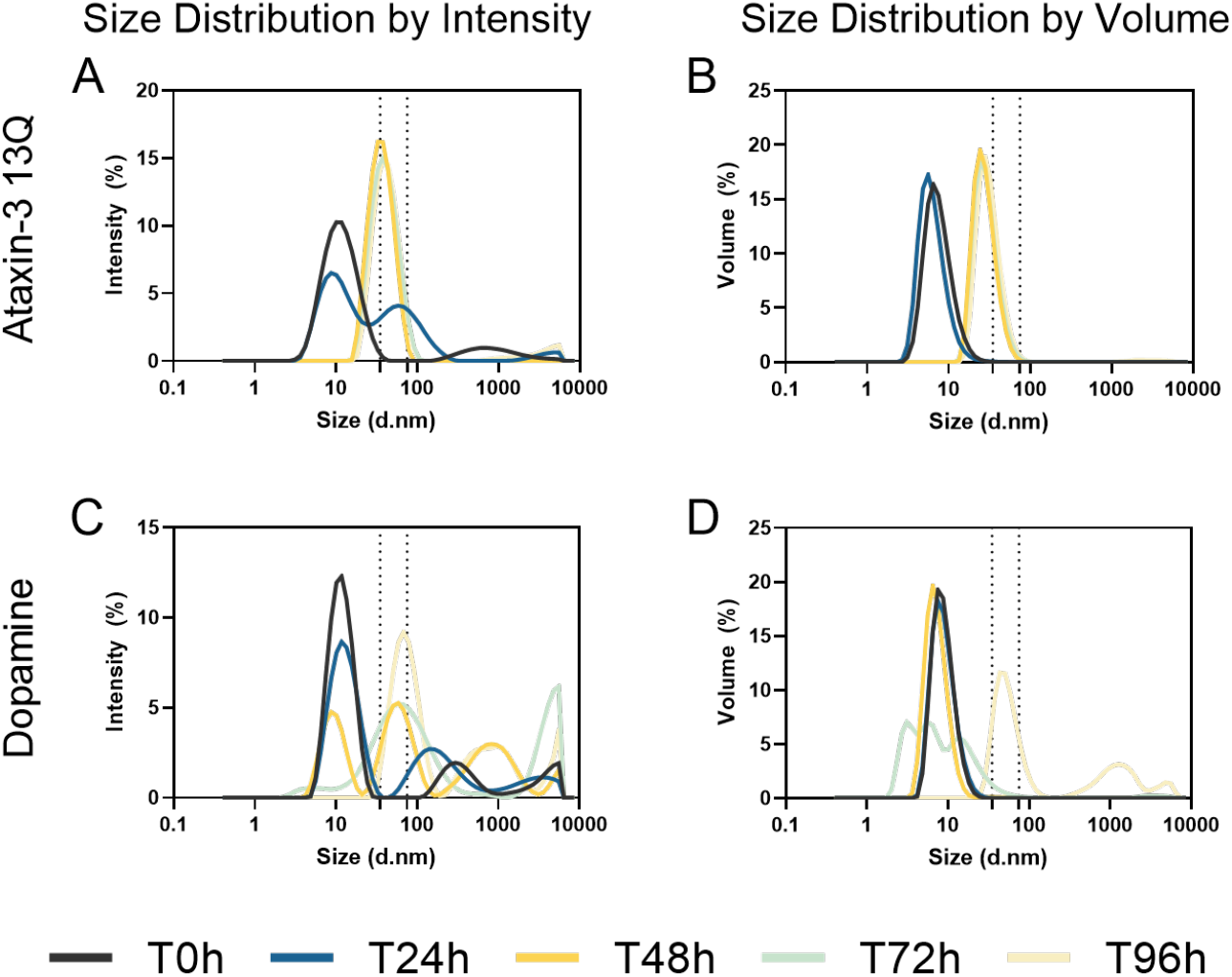
Effect of dopamine hydrochloride on Atx3 13Q aggregation analyzed by DLS. Size distributions measured in the absence (A, B) and presence (C, D) of 5 μM dopamine hydrochloride. Left (A, C) and right (B, D) panels refer to size distributions by intensity and volume, respectively. Solid lines: mean of three independent replicates for each tested condition. Dashed lines: guides indicating particle sizes of 35 and 75 nm in the size distributions.

**Figure 5.**
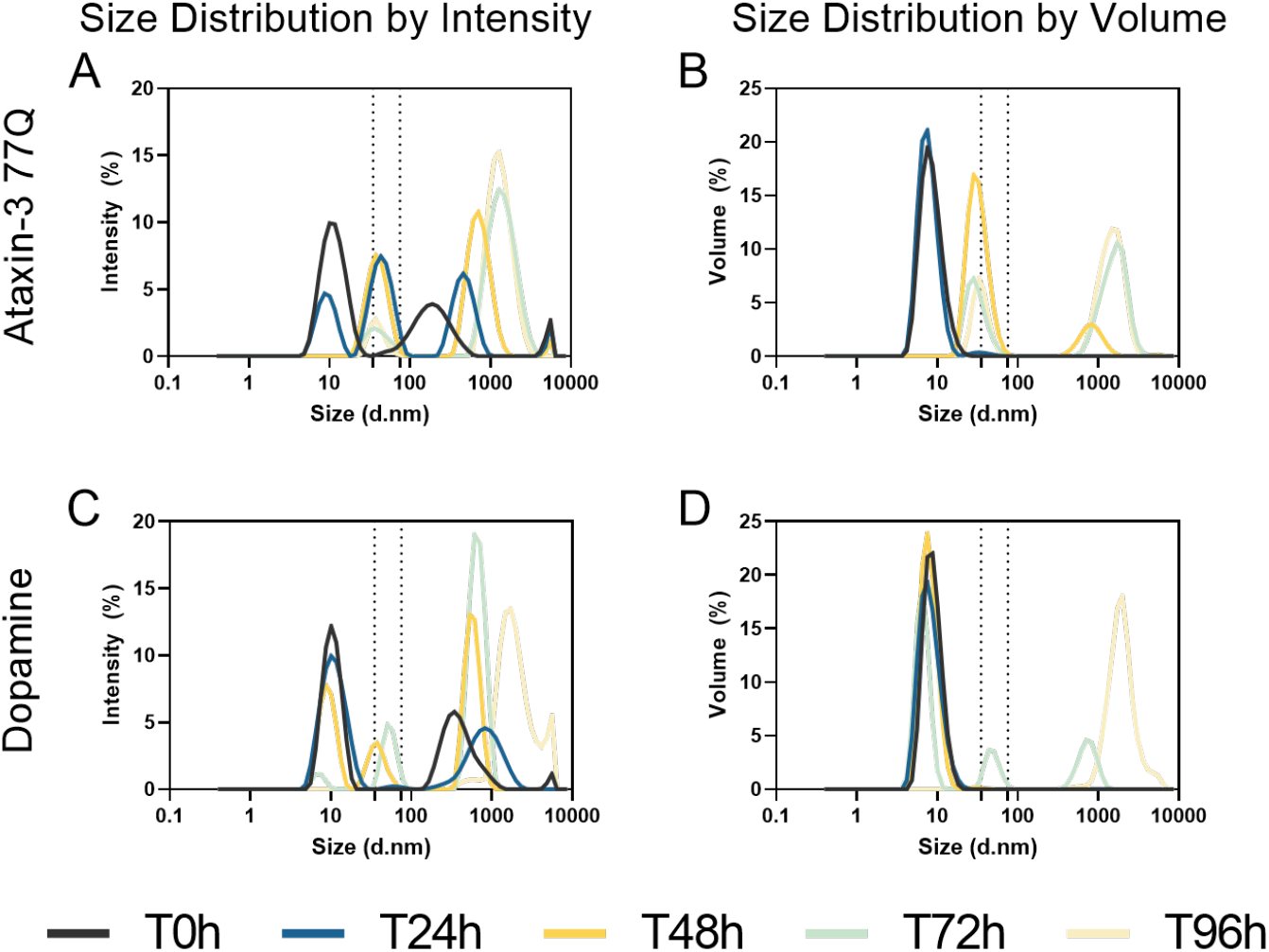
Effect of dopamine hydrochloride on Atx3 77Q aggregation analyzed by DLS. Size distributions measured in the absence (A, B) and presence (C, D) of 5 μM dopamine hydrochloride. Left (A, C) and right (B, D) panels refer to size distributions by intensity and volume, respectively. Solid lines: mean of three independent replicates for each tested condition. Dashed lines: guides indicating particle sizes of 35 and 75 nm in the size distributions.

Larger aggregates produced at slower aggregation rates were also observed for tolcapone (Figure S2) ciclopirox ethanolamine (Figure S4) and pentetic acid (Figure S6). As previously demonstrated for Atx3 13Q [24], the average fibril size reflects the relative importance of the 1ry and 2ry nucleation steps, with smaller fibrils reflecting predominant (i.e., faster) 2ry nucleation. Concerning Atx3 77Q aggregation, an additional population of species larger than 100 nm is identified from the beginning of incubation if intensity distributions are analyzed (Figures 5A and 5C). With the further formation of large aggregates, most likely mature fibrils [21], this population becomes visible in the volume distributions as well, first in the control (Figure 5B) and then in the test compound (Figure 5D) experiment. Similarly to Atx3 13Q, 5 μM dopamine delays the 2ry nucleation step and monomer depletion as indicated by the later appearance of >35 nm peaks and disappearance of the ~10 nm peak. Although in lesser amounts than in the control, Atx3 77Q large aggregates can be identified in the volume distributions after 72 h incubation in the presence of dopamine (Figure 5D) and tolcapone (Figure S3). A similar analysis for ciclopirox ethanolamine (Figure S5) and pentetic acid (Figure S7) does not reveal the presence of detectable amounts of large aggregates. Together with the kinetic analysis of the *t*_lag_ and *ν*_max_ measurables (Figures 3A and 3C), our DLS results support a common inhibition mechanism where the formation of new fibrils by 2ry nucleation is markedly delayed. Comparatively, the steps of 1ry nucleation and fibril elongation are much less affected by the presence of the inhibitors. The step of Atx3 77Q fibril maturation is also delayed but, in this case, the inhibitory effect is more pronounced for ciclopirox ethanolamine and pentetic acid.

### Dopamine delays the formation of precursor oligomers and mature fibrils

By preventing the early formation of 2ry fibrils common to the aggregation pathways of Atx3 13Q and Atx3 77Q, we expect that the formation of disease-characteristic mature fibrils of Atx3 77Q is also prevented. To validate this mechanism, complementary experiments were performed using SEC and TEM to monitor Atx3 and 6F). As quantitatively illustrated by the relative peak weights of HWM oligomers and Atx3 monomers, most of the protein remained monomeric in the presence of 1 and 5 μM dopamine after 68 hours incubation, while in the control experiments both non-expanded and expanded Atx3 were fully aggregated in less than 44 hours. Similar inhibition of HMW oligomer formation aggregation over time. Unlike DLS, SEC analysis is focused only on the populations of soluble protein and oligomers that are loaded into the SEC column after sample filtration. This allowed us to confirm that the formation of high molecular weight (HMW) oligomers assembled during the early stages of aggregation is delayed in the case of Atx3 13Q (Figures 6A, 6C, and 6E), but also in the case of Atx3 77Q (Figures 6B, 6D was observed for 1 and 5 μM tolcapone (Figure S8) and by 2.5 μM pentetic acid (Figure S9). Although the formation of HMW oligomers of Atx3 was delayed by 0.5 and 2.5 μM ciclopirox ethanolamine, this compound did not prevent the ultimate conversion of most monomers into high-order aggregates (Figure S10).

**Figure 6.**
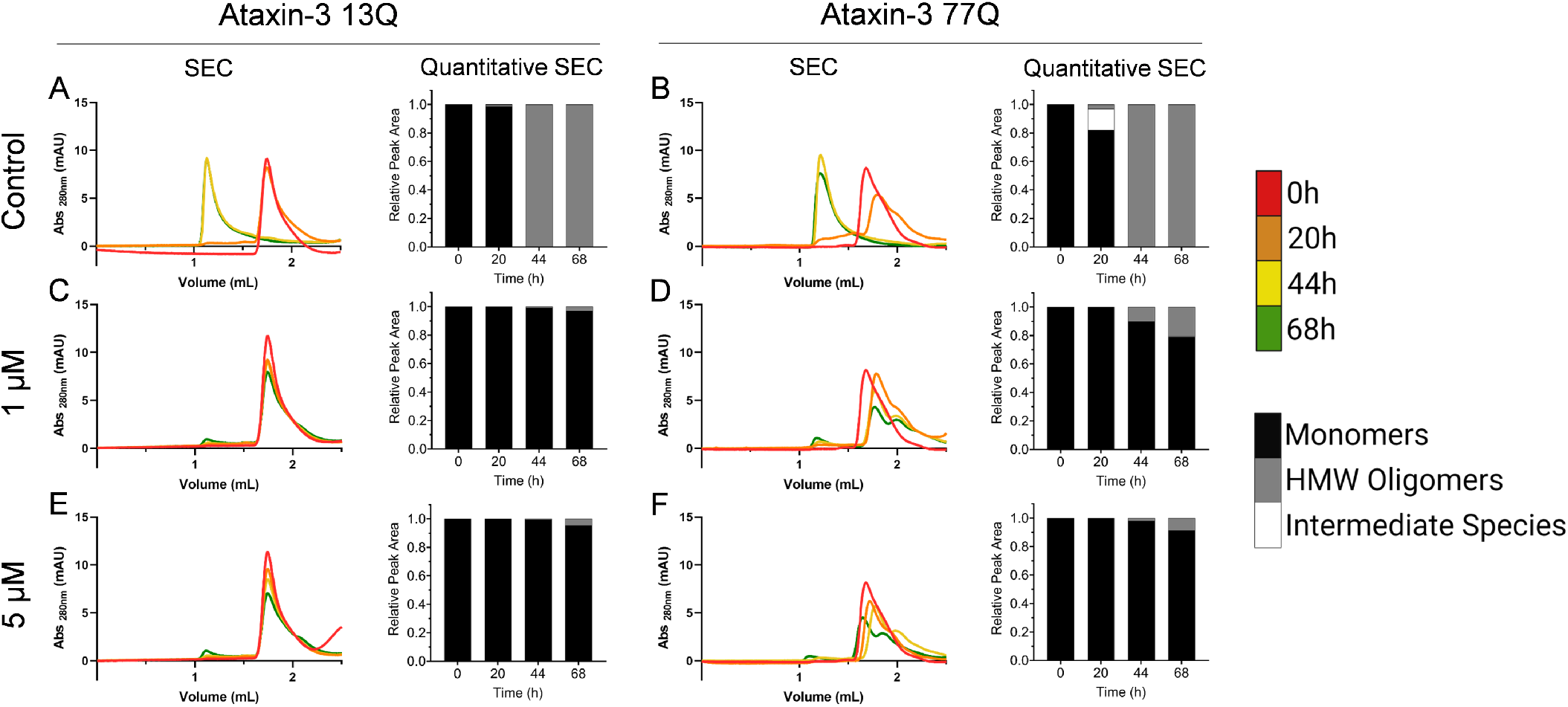
Effect of dopamine hydrochloride on Atx3 aggregation analyzed by SEC. Time-course chromatograms during the aggregation of Atx3 13Q (A, C and E) and Atx3 77Q (B, D and F) in the absence (A and B) and presence of 1 μM (C and D) and 5 μM (E and F) dopamine hydrochloride. Different colors: different incubation times as indicated at the color scale. On the right side of each chromatogram is the corresponding quantitative SEC analysis, which is expressed in relative peak area units of monomer (black), high molecular weight oligomers (grey), and intermediate species (white).

To further evaluate whether the formation of Atx3 aggregates, in particular the appearance of Atx3 77Q mature fibrils, is inhibited by dopamine, the same aliquots used for SEC analysis were observed without prior filtration by TEM. Using this technique we could identify oligomeric structures, small fibrils and, in the case of Atx3 77Q, large mature fibrils. The appearance of all types of aggregates was markedly delayed in the presence of 1 and 5 μM dopamine (Figure 7). Although mature fibrils of Atx3 77Q were not totally suppressed by dopamine, their quantity was visibly reduced in the presence of this inhibitor. A similar effect was identified for the studied concentrations of tolcapone (Figure S11). A stronger reduction in the number of mature fibrils was observed for pentetic acid (Figure S12) in agreement with the results obtained by DLS (Figure S7D). Ciclopirox ethanolamine (Figure S13) seems to have a weaker inhibitory effect on Atx3 77Q assembly than the one indicated by DLS measurements (Figure S5D). This difference is understandable given the non-quantitative information provide by TEM and the higher compound concentrations used during DLS measurements. Overall, the complementary techniques used to characterize the inhibition mechanisms at play during Atx3 aggregation confirm that the early steps of secondary nucleation, as well as the later steps of Atx3 77Q fibril maturation, are prevented or delayed in the presence of dopamine and 3 other drug-repurposing compounds.

**Figure 7.**
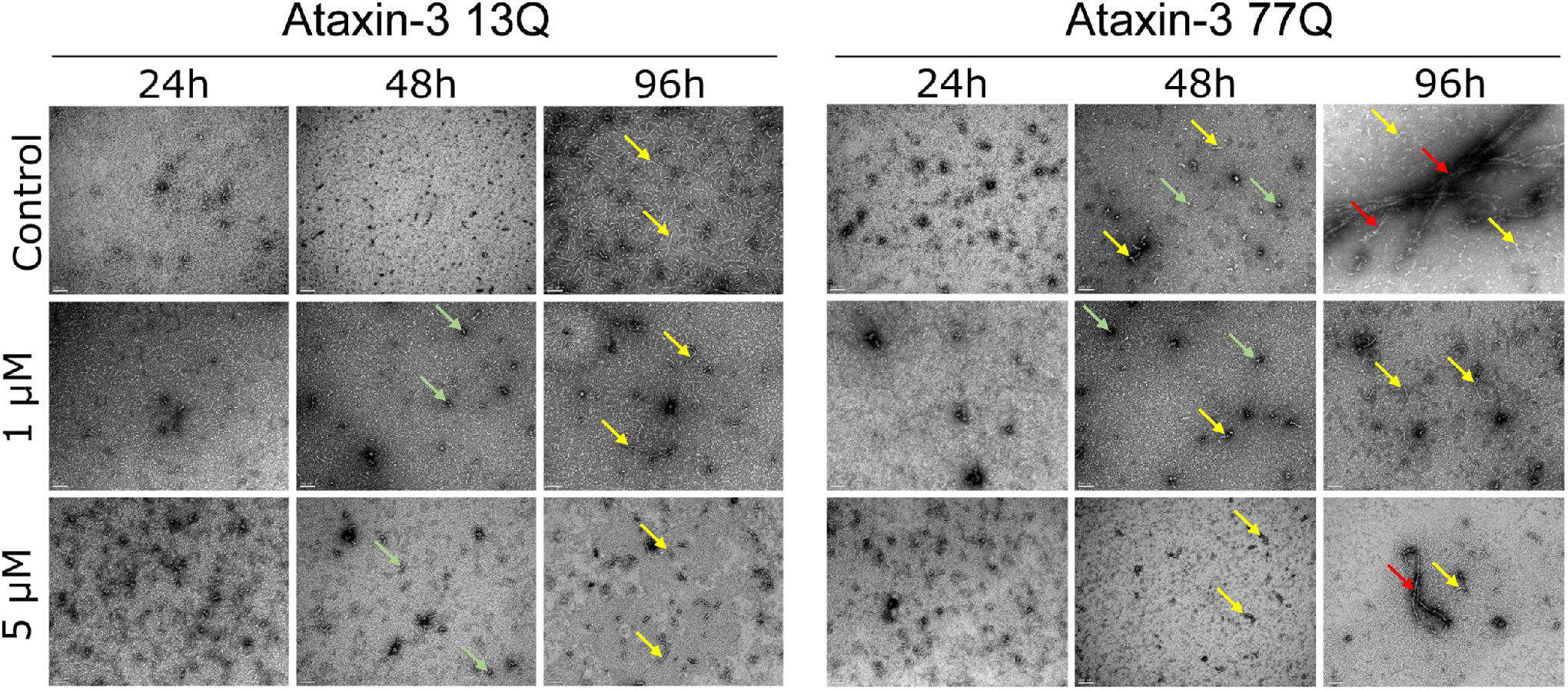
Effect of dopamine hydrochloride on Atx3 aggregation analyzed by TEM. TEM images of negatively stained Atx3 13Q (series on the left) and Atx3 77Q (series on the right) incubated over different periods (24, 48 and 96 h) in the absence (top series) and presence of 1 μM (middle series) and 5 μM (bottom series) of dopamine hydrochloride. Scale bars correspond to 100 nm. Examples of HMW oligomers, small fibrils and mature fibrils are highlighted by green, yellow and red arrows, respectively.

## Discussion

The landscape of coexisting Atx3 species includes soluble oligomers, ThT-positive fibrils formed either by primary and secondary nucleation, and, in the case of polyQ-expanded Atx3, SDS-resistant mature fibrils formed by the self-assembly of the precursor fibrils. In this drug repurposing study, we aimed at identifying kinetic inhibitors of the earlier steps of protein aggregation so that new therapeutic interventions targetting NNI accumulation can be eventually designed. While it is not clear which of the species actively participate in the cascade of MJD pathogenesis, it is expected that the kinetic inhibition of the nucleation steps would delay the formation of (i) disease-characteristic straight mature fibrils, associated with cytotoxicity in Huntington’s disease [2,34,35], and (ii) seed-competent fibrils/protofibrils that are likely involved in interneuronal disease spreading [36]. To be translated into clinical practice, such a preventive treatment should be safely tolerated in a chronic medication regime started during the early or pre-symptomatic stages of the disease. Given this limitation, the discovery of dopamine as an effective inhibitor of Atx3 aggregation is of particular interest as it opens the possibility of using dopaminergic drugs in MJD, not only to treat parkinsonian symptoms but also as a long-term, low-dose medication preventing disease progression. The neurotransmitter dopamine belongs to the chemical family of catecholamines that are known inhibitors of other amyloidogenic proteins and peptides such as α-synuclein, amyloid-ß peptide, and human islet amyloid polypeptide [37–39]. Differently from those cases, Atx3 aggregation into ThT-positive fibrils is mediated by a protein domain with a known and well-characterized structure – the Josephin domain. In support of a nonspecific anti-amyloid activity of dopamine, the results obtained by the thermal shift assay (Figure S1) indeed suggest no particular stabilization or destabilization of the Atx3 monomer resulting from interactions with this catecholamine. Uncovering the inhibition mechanism of amyloid aggregation by catecholamines is difficult due to the auto-oxidation of catechols to quinones and other sub-products [40]. Additionally, in the specific case of α-synuclein, a non-amyloidogenic pathway is elicited by the presence of dopamine involving the formation of covalent adducts and small protein oligomers [39,41]. Our analysis of Atx3 aggregation using gel filtration chromatography suggests that the population of small oligomers of Atx3 detectable in SEC chromatograms is greatly reduced by the presence of dopamine (Figure 6). Moreover, the observation of non-filtered samples by TEM does not indicate the presence of additional populations of protein aggregates generated from the interaction with our inhibitors (Figure 7). As on previous occasions [40], the effects of dopamine can hardly be separated from those of dopamine-oxidation products such as dopamine *o*-quinone and 5,6-indolequinone. It seems clear, however, that the early steps of Atx3 aggregation are kinetically inhibited by the presence of sub-micromolar concentrations of dopamine (Figures 2 and 3) as manifested by the marked reduction of HMW oligomers, fibrils/protofibrils and mature fibril content (Figures 4, 6 and 7).

Also included in our selection of effective inhibitors is tolcapone, which, similarly to dopamine, has an anti-amyloid activity demonstrated for other polypeptides like α-synuclein, amyloid-β peptide, and transthyretin [31,42]. Pentetic acid, on the other hand, significantly delayed the self-assembly of Atx3 monomers for lower compound concentrations than dopamine or tolcapone. From a previous screening targeting the deubiquitinase activity of Atx3 [43], we know that none of the selected compounds was identified as affecting the enzymatic activity of this protein. The main reason why dopamine is here highlighted as the most promising candidate for drug repurposing in MJD is the existence of strong preclinical and clinical evidence showing, in the context of Parkinson’s disease (PD), that dopamine brain levels can be safely increased through the administration dopaminergic drugs such as levodopa (levodihydoxyphenylalanine, a.k.a. L-dopa) and dopamine agonists [44]. Since its introduction in the late 1960s [45], levodopa has been the gold standard for the therapy of motor symptoms such as tremor, rigidity and bradykinesia related to dopamine deficiency [46]. Measurements of the dopamine concentration in the extracellular fluid of rodent brains before and after the stimulation of dopaminergic terminals point to nanomolar and micromolar levels, respectively [47,48], which are in line with later estimates of striatal dopamine fluctuations in humans [49]. Restoration of this neurotransmitter is possible through the administration of its metabolic precursor levodopa, which is a natural substrate for the enzyme aromatic-amino-acid decarboxylase catalyzing the formation of dopamine [40], but also by directly acting on dopamine receptors through the administration of dopamine agonists [50]. Levodopa is typically dosed 3 times daily to start but, as PD progresses, higher and more frequent doses are required to account for the decreasing ability to store dopamine excess and the decreasing duration of response to dopaminergic medication [44]. Low doses (300-600 mg/day) of levodopa can effectively control PD symptoms [51], while several studies by Japanese child neurologists confirm the positive effect of very low doses (0.5-1 mg/kg/day) on pediatric neurological disorders [52]. The benefits provided by levodopa to millions of PD patients are not without adverse effects such as nausea and vomiting as a consequence of the overproduction of dopamine by the peripheral metabolism [46]. These effects can be prevented by including BBB-impermeable decarboxylase inhibitors such as carbidopa in the levodopa formulation. High doses of levodopa are associated with dyskinesia symptoms, i.e., fast, unpredictable, and involuntary movements manifested by PD patients (usually) years after the treatment is initiated [44,46]. According to recent clinical evidence, dyskinesia development is a function of disease duration rather than cumulative levodopa exposure, and therefore postponing levodopa treatment to avoid its side effects is discouraged [50]. More efficient delivery systems than the short-acting oral formulations improve dyskinesia symptoms, which can also be managed with extended-release formulations of amantadine [44,46,50].

In our protein aggregation assays, the formation of Atx3 fibrils is efficiently inhibited for dopamine concentrations that are within the physiological boundaries estimated from studies in rodents and humans. Interestingly, the antipsychotic aripirazole, shown to decrease the levels of polyQ-expanded Atx3 in the brain of MJD transgenic mice without directly interfering with Atx3 fibril assembly *in vitro* [53], is a partial agonist of dopamine D2 receptors in mammals [54]. Recently, it has been demonstrated that dopamine D2-like receptors are partly responsible for aripiprazole-mediated improvement motor dysfunction in a *C. elegans* MJD model [55]. A therapeutic strategy relying on low doses of dopaminergic drugs to prevent NNI accumulation would counteract a possible impairment of the nigrostriatal dopaminergic system associated with MJD [56], a feature that was previously identified in both symptomatic and asymptomatic MJD disease gene carriers using single-photon emission computed tomography [57,58]. Moreover, transcranial brain sonography studies reveal a significantly higher frequency of substantia nigra hyper echogenicity in MJD patients [59], while it is known from the early neuropathological descriptions of MJD that the substantia nigra, together with the dentate nucleus of the cerebellum, are almost invariably the main targets of the disease [1]. Although extrapyramidal signs such as dystonia and rigidity can be masked in patients with dominant spinocerebellar ataxia [10,57], subtypes I and III of MJD feature extrapyramidal signs and subtype IV features levodopa-responsive parkinsonism [9,10,60,61]. Finally, an interesting case report was documented in 2021 about a patient presenting signs indistinguishable from Parkinson’s disease who was later diagnosed with MJD; it was not until ten years of treatment with levodopa and other dopamine agonists the patient developed the classical MJD symptoms, cerebellar ataxia and pyramidal signs [11].

Knowing the impact of candidate therapies on neurodegenerative diseases has been hindered by the lack of clear clinical endpoints, biomarkers or surrogate markers besides the quantitative neurological scales of disease progression. Distinctively from using dopaminergic drugs to treat parkinsonian symptoms in a minority of MJD patients, halting or slowing down MJD progression is, in principle, oriented to all affected populations and would require prolonged clinical trials to validate the drugs’ new use. Even under accelerated drug approval pathways, these clinical trials are associated with a high financial risk for pharmaceutical sponsorship unless additional investment from the public sector is involved [26,62]. Because MJD is a rare genetic disorder whose clinical features correlate with the length of the CAG repeat [63], preventive trials can alternatively be designed to follow drug-treated presymptomatic individuals to the clinical onset and compare their response with that of historical controls [62]. Successful stories such as the treatment from the childhood of familial hypercholesterolemia are, in this regard, inspirational for future preventive therapies stemming from bench research [64].

## Conclusions

Our *in vitro* results show that physiological concentrations of dopamine are sufficient to significantly prevent or delay the aggregation of Atx3 and the ultimate formation of mature fibrils characteristic of Atx3 with pathological polyQ tract sizes. Other potent inhibitors identified in our drug repurposing study show less favourable safety and pharmacokinetic profiles than well-known dopamine precursors and dopamine agonists used in varying doses to treat PD symptoms. Considering our results, the available pharmacological data, and the neuropathological evidence for the involvement of the nigrostriatal dopaminergic system in MJD pathogenesis, we purpose that low to very low doses of dopaminergic drugs should be further investigated as a means to prevent NNI accumulation by keeping balanced levels of dopamine in the brains of MJD gene carriers. To be convincingly tested, such a strategy will require randomized clinical trials showing clear disease-modifying effects or that the clinical onset of MJD can be prevented or delayed by dopaminergic drugs.

## Acknowledgements

This work is part of a project that has received funding from the European Union’s Horizon 2020 research and innovation programme under grant agreement No. 952334 (PhasAGE). This research was financed by FEDER—Fundo Europeu de Desenvolvimento Regional funds through the COMPETE 2020—Operational Programme for Competitiveness and Internationalization (POCI), Portugal 2020, and by Portuguese funds through FCT—Fundação para a Ciência e a Tecnologia/ Ministério da Ciência, Tecnologia e Ensino Superior (FCT/MCTES) in the framework of projects PTDC/QUI-COL/2444/2021, POCI-01-0145-FEDER-031323 (PTDC/MED-FAR/31323/2017), and POCI-01-0145-FEDER-007274 (“Institute for Research and Innovation in Health Sciences”).

