## Supplementary Figures for "Drug repurposing of dopaminergic drugs to inhibit Ataxin-3 aggregation"

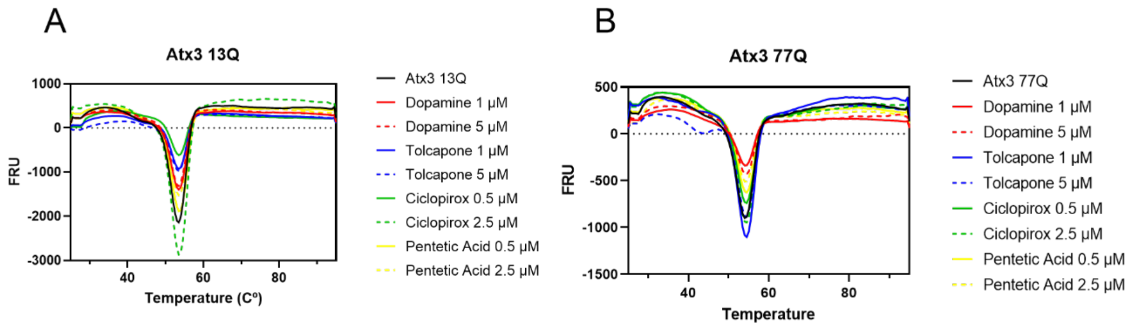


**Figure S1**. Thermal shift analysis reveals no high-affinity interactions. Fluorescence signal measured for increasing temperatures of (A) Atx3 13Q and (B) Atx3 77Q incubated with each compound on a buffer containing SYPRO Orange dye.

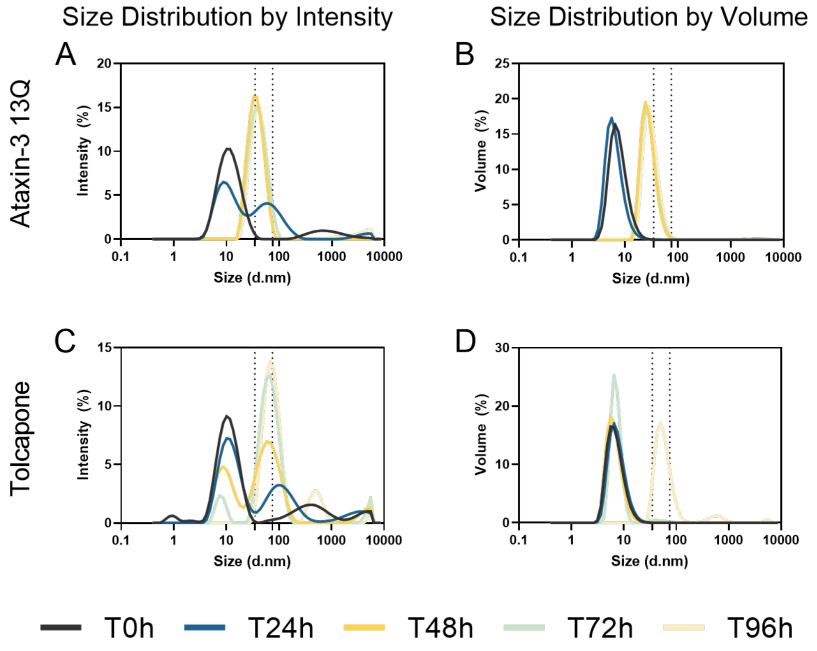


**Figure S2**. Effect of tolcapone on Atx3 13Q aggregation analyzed by DLS. Size distributions measured in the absence (A, B) and presence (C, D) of 5 µM tolcapone. Left (A, C) and right (B, D) panels refer to size distributions by intensity and volume, respectively. Solid lines: mean of three independent replicates for each tested condition. Dashed lines: guides indicating particle sizes of 35 and 75 nm in the size distributions.

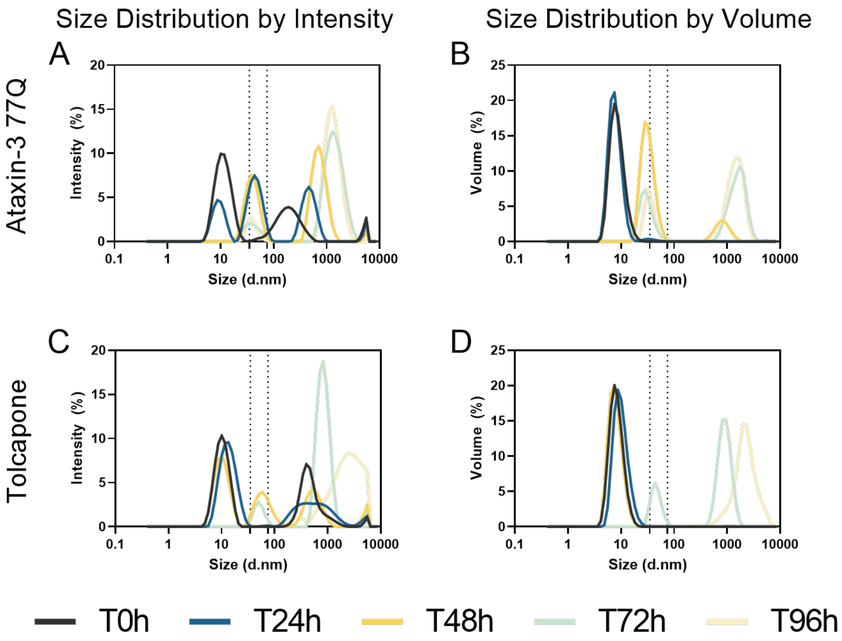


**Figure S3**. Effect of tolcapone on Atx3 77Q aggregation analyzed by DLS. Size distributions measured in the absence (A, B) and presence (C, D) of 5 µM tolcapone. Left (A, C) and right (B, D) panels refer to size distributions by intensity and volume, respectively. Solid lines: mean of three independent replicates for each tested condition. Dashed lines: guides indicating particle sizes of 35 and 75 nm in the size distributions.

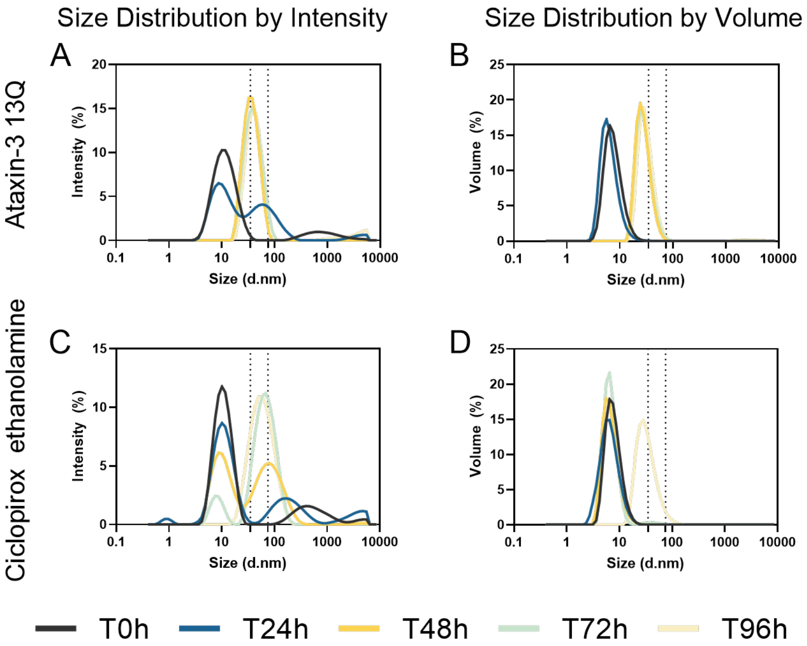


**Figure S4**. Effect of ciclopirox ethanolamine on Atx3 13Q aggregation analyzed by DLS. Size distributions measured in the absence (A, B) and presence (C, D) of 5 µM ciclopirox ethanolamine. Left (A, C) and right (B, D) panels refer to size distributions by intensity and volume, respectively. Solid lines: mean of three independent replicates for each tested condition. Dashed lines: guides indicating particle sizes of 35 and 75 nm in the size distributions.

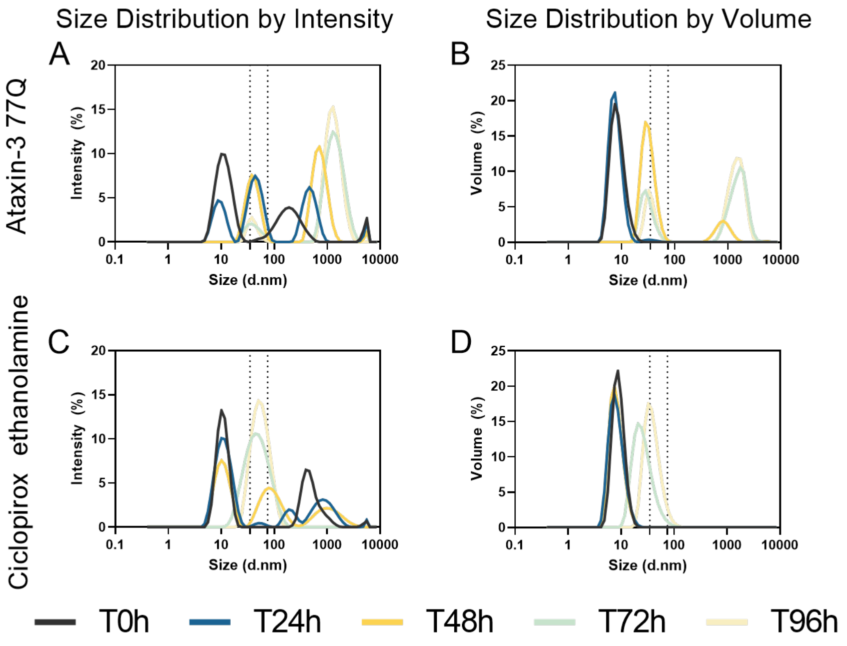


**Figure S5**. Effect of ciclopirox ethanolamine on Atx3 77Q aggregation analyzed by DLS. Size distributions measured in the absence (A, B) and presence (C, D) of 5 µM ciclopirox ethanolamine. Left (A, C) and right (B, D) panels refer to size distributions by intensity and volume, respectively. Solid lines: mean of three independent replicates for each tested condition. Dashed lines: guides indicating particle sizes of 35 and 75 nm in the size distributions.

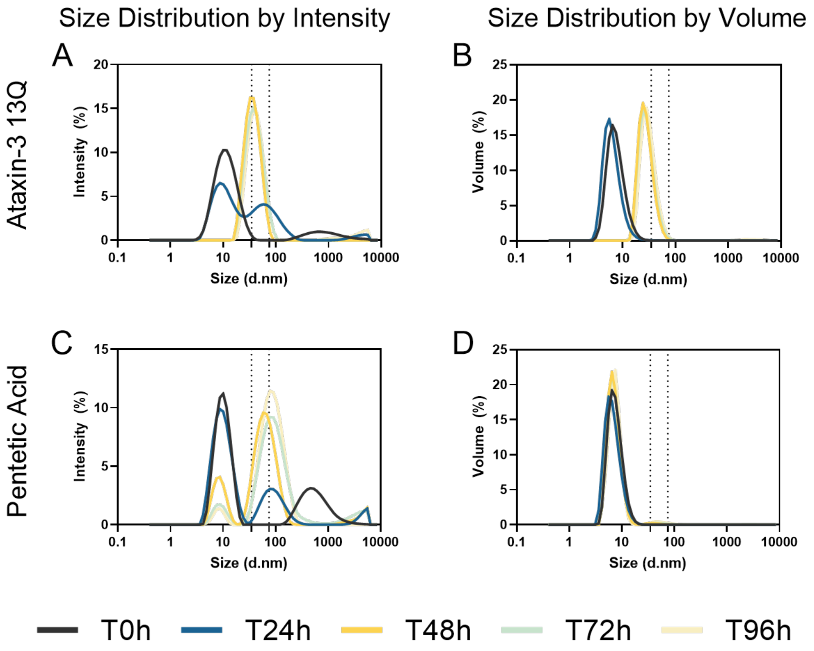


**Figure S6**. Effect of pentetic acid on Atx3 13Q aggregation analyzed by DLS. Size distributions measured in the absence (A, B) and presence (C, D) of 5 µM pentetic acid. Left (A, C) and right (B, D) panels refer to size distributions by intensity and volume, respectively. Solid lines: mean of three independent replicates for each tested condition. Dashed lines: guides indicating particle sizes of 35 and 75 nm in the size distributions.

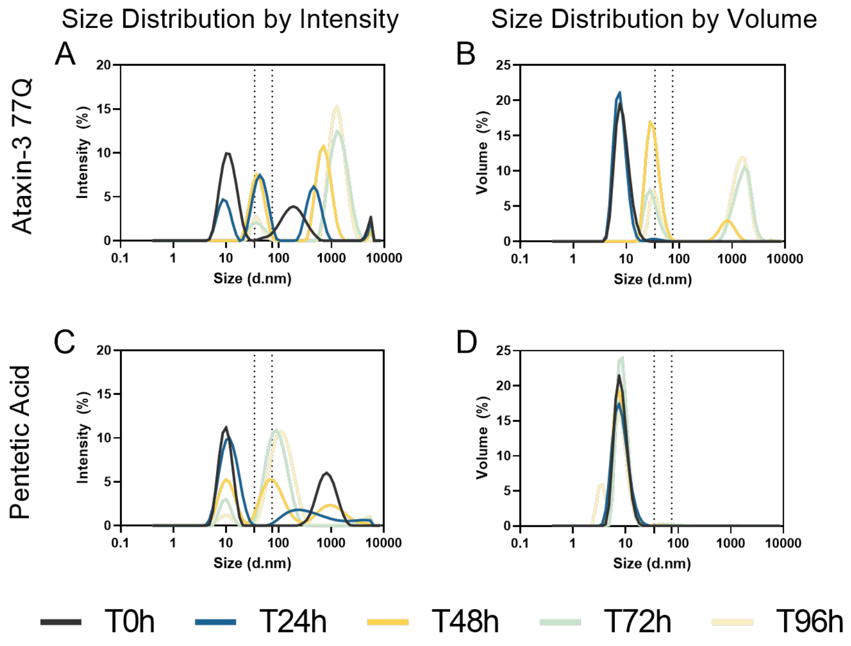


**Figure S7**. Effect of pentetic acid on Atx3 77Q aggregation analyzed by DLS. Size distributions measured in the absence (A, B) and presence (C, D) of 5 µM pentetic acid. Left (A, C) and right (B, D) panels refer to size distributions by intensity and volume, respectively. Solid lines: mean of three independent replicates for each tested condition. Dashed lines: guides indicating particle sizes of 35 and 75 nm in the size distributions.

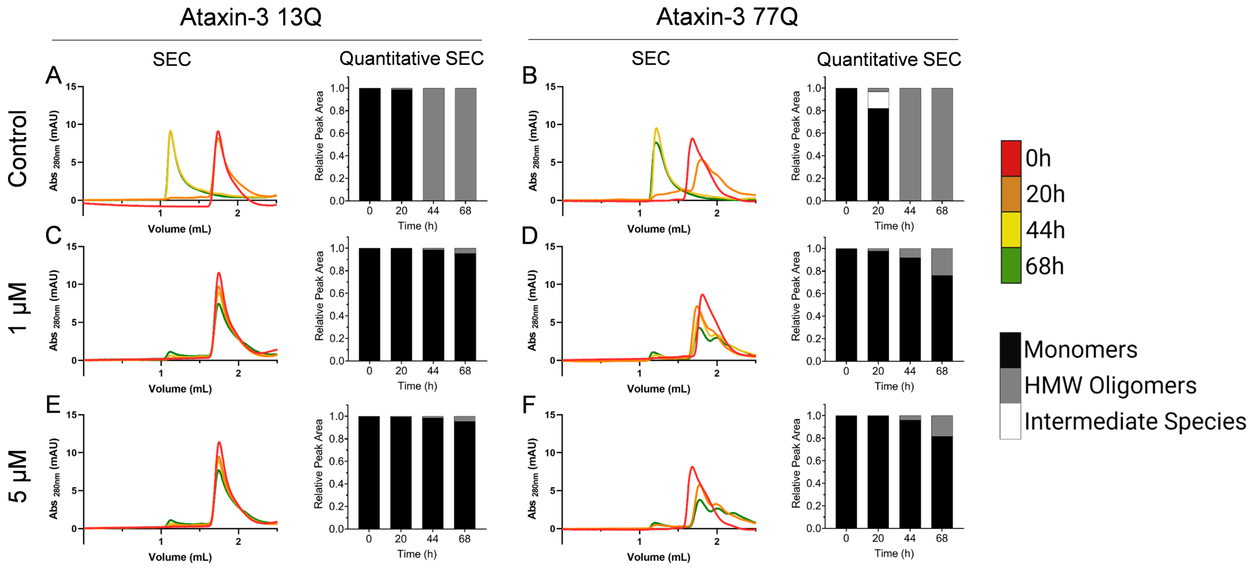


**Figure S8**. Effect of tolcapone on Atx3 aggregation analyzed by SEC. Time-course chromatograms during the aggregation of Atx3 13Q (A, C and E) and Atx3 77Q (B, D and F) in the absence (A and B) and presence of 1 µM (C and D) and 5 µM (E and F) tolcapone. Different colors: different incubation times as indicated at the color scale. On the right side of each chromatogram is the corresponding quantitative SEC analysis, which is expressed in relative peak area units of monomer (black), high molecular weight oligomers (grey), and intermediate species (white).

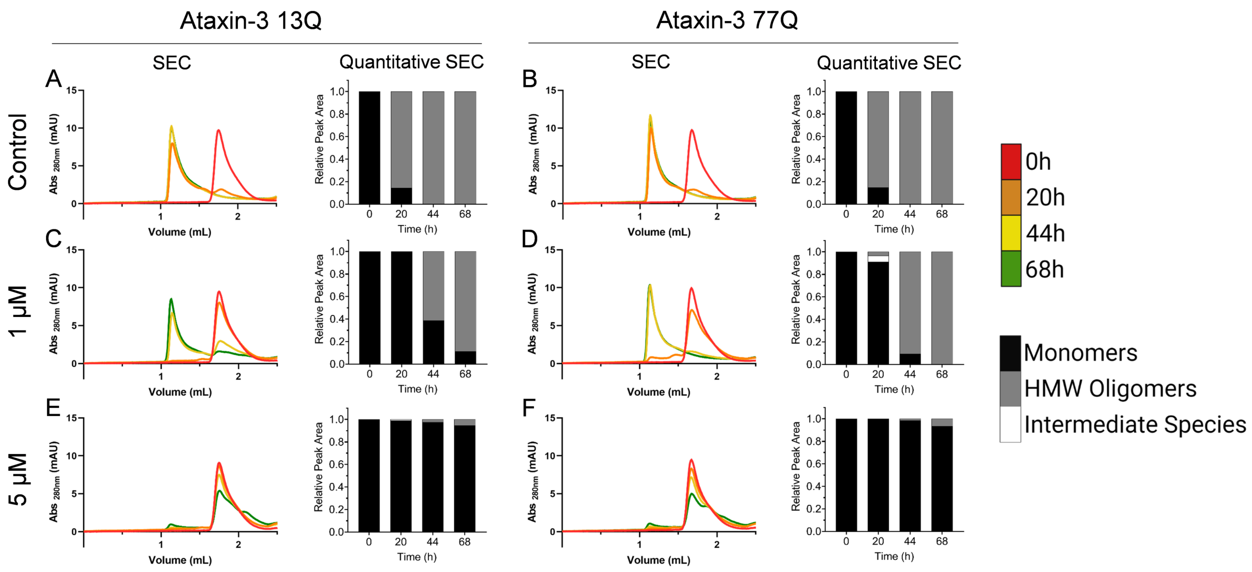


**Figure S9**. Effect of pentetic acid on Atx3 aggregation analyzed by SEC. Time-course chromatograms during the aggregation of Atx3 13Q (A, C and E) and Atx3 77Q (B, D and F) in the absence (A and B) and presence of 0.5 µM (C and D) and 2.5 µM (E and F) pentetic acid. Different colors: different incubation times as indicated at the color scale. On the right side of each chromatogram is the corresponding quantitative SEC analysis, which is expressed in relative peak area units of monomer (black), high molecular weight oligomers (grey), and intermediate species (white).

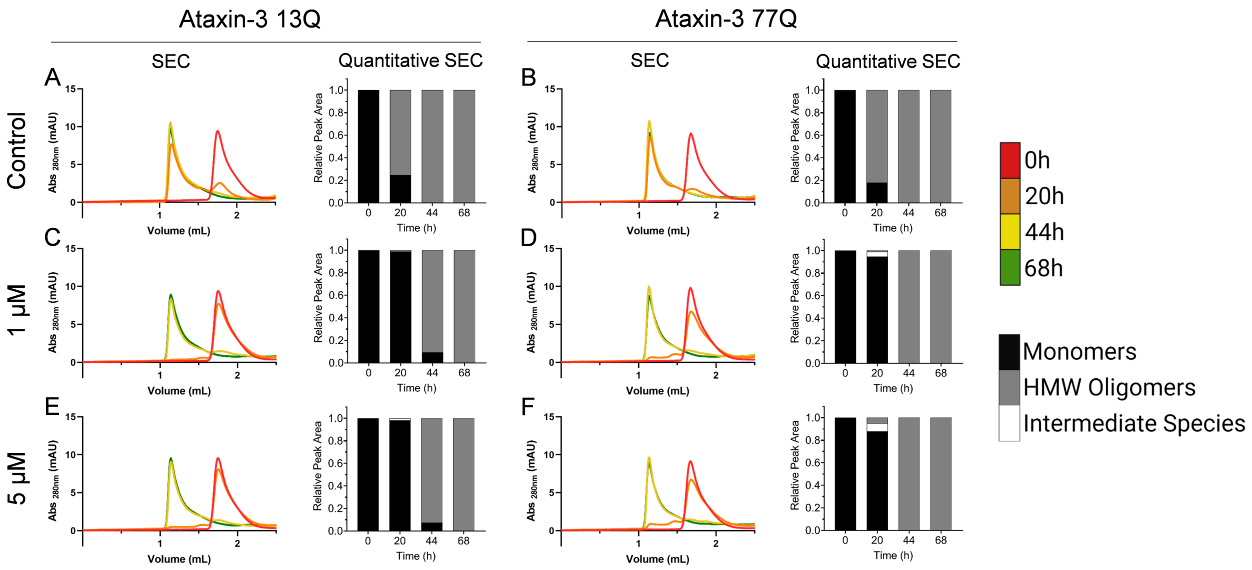


**Figure S10**. Effect of ciclopirox ethanolamine on Atx3 aggregation analyzed by SEC. Time-course chromatograms during the aggregation of Atx3 13Q (A, C and E) and Atx3 77Q (B, D and F) in the absence (A and B) and presence of 0.5 µM (C and D) and 2.5 µM (E and F) ciclopirox ethanolamine. Different colors: different incubation times as indicated at the color scale. On the right side of each chromatogram is the corresponding quantitative SEC analysis, which is expressed in relative peak area units of monomer (black), high molecular weight oligomers (grey), and intermediate species (white).

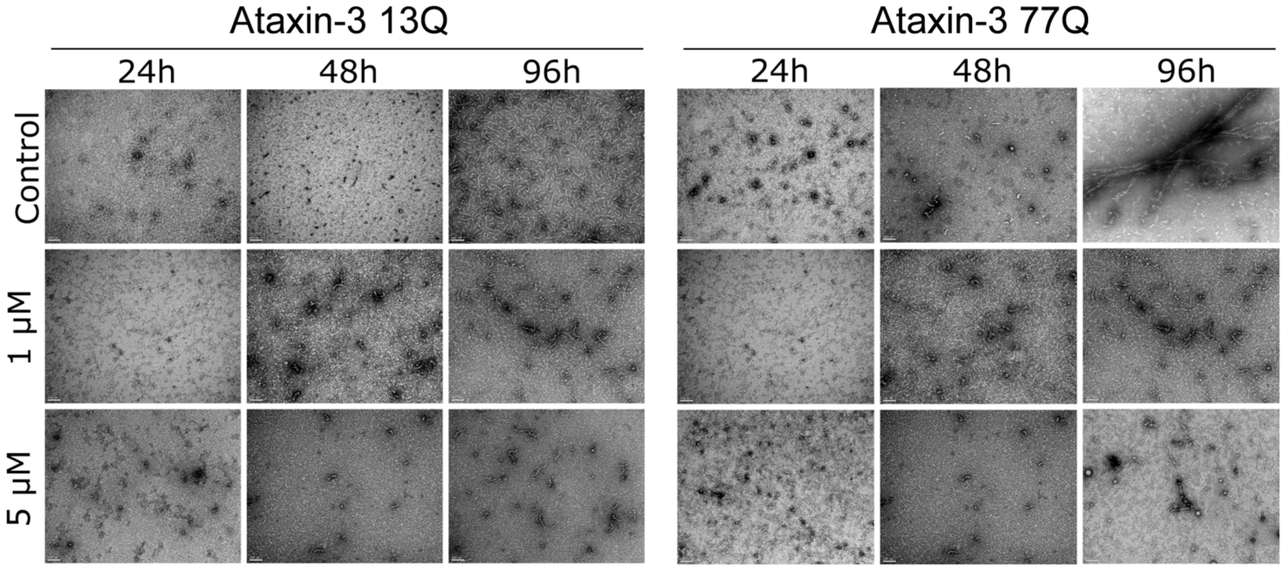


**Figure S11**. Effect of tolcapone on Atx3 aggregation analyzed by TEM. TEM images of negatively stained Atx3 13Q (series on the left) and Atx3 77Q (series on the right) incubated over different periods (24, 48 and 96 h) in the absence (top series) and presence of 1 µM (middle series) and 5 µM (bottom series) of tolcapone. Scale bars correspond to 100 nm.

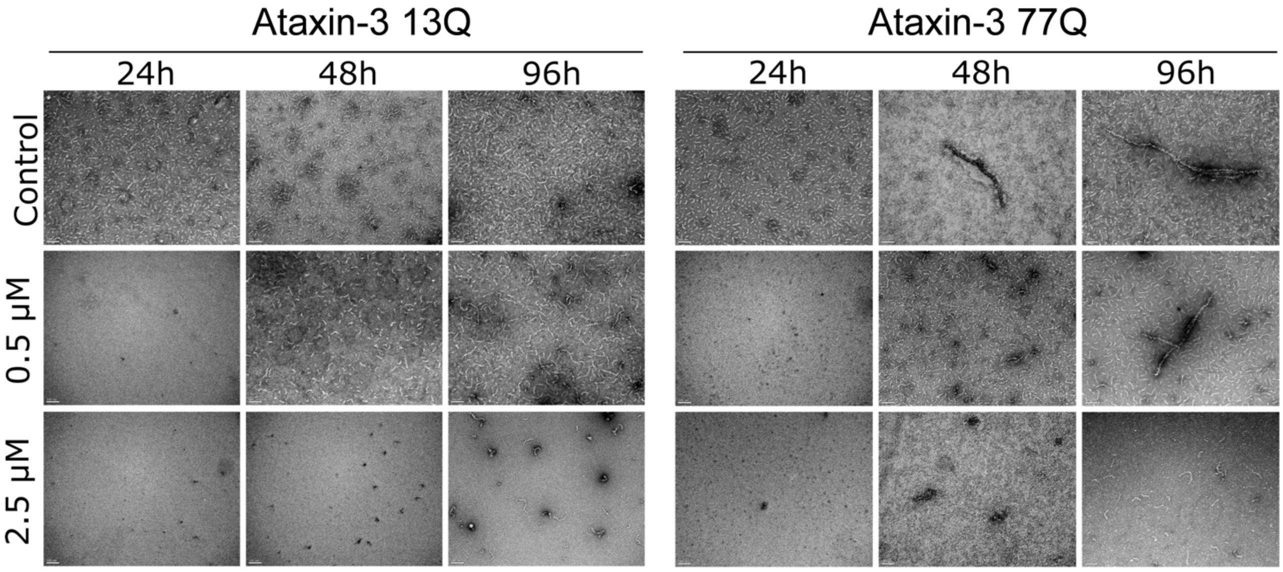


**Figure S12**. Effect of pentetic acid on Atx3 aggregation analyzed by TEM. TEM images of negatively stained Atx3 13Q (series on the left) and Atx3 77Q (series on the right) incubated over different periods (24, 48 and 96 h) in the absence (top series) and presence of 0.5 µM (middle series) and 2.5 µM (bottom series) of pentetic acid. Scale bars correspond to 100 nm.

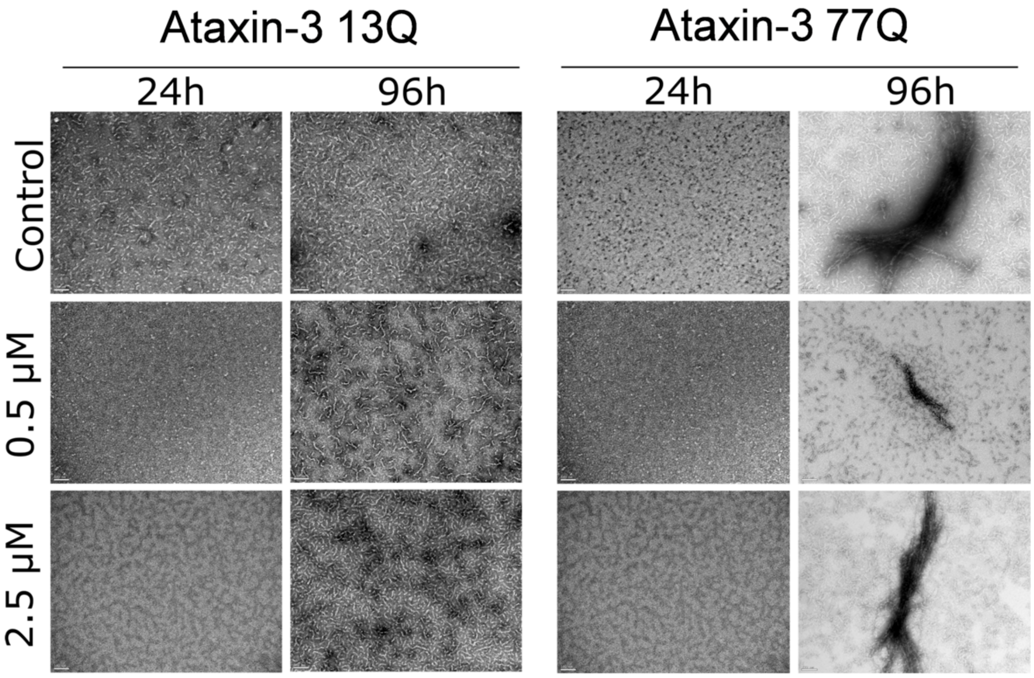


**Figure S13**. Effect of ciclopirox ethanolamine on Atx3 aggregation analyzed by TEM. TEM images of negatively stained Atx3 13Q (series on the left) and Atx3 77Q (series on the right) incubated over different periods (24 and 96 h) in the absence (top series) and presence of 0.5 μM (middle series) and 2.5 μM (bottom series) of ciclopirox ethanolamine. Scale bars correspond to 100 nm.
